# TMPRSS2 and TMPRSS4 mediate SARS-CoV-2 infection of human small intestinal enterocytes

**DOI:** 10.1101/2020.04.21.054015

**Authors:** Ruochen Zang, Maria F.G. Castro, Broc T. McCune, Qiru Zeng, Paul W. Rothlauf, Naomi M. Sonnek, Zhuoming Liu, Kevin F. Brulois, Xin Wang, Harry B. Greenberg, Michael S. Diamond, Matthew A. Ciorba, Sean P.J. Whelan, Siyuan Ding

## Abstract

Both gastrointestinal symptoms and fecal shedding of SARS-CoV-2 RNA have been frequently observed in COVID-19 patients. However, whether SARS-CoV-2 replicate in the human intestine and its clinical relevance to potential fecal-oral transmission remain unclear. Here, we demonstrate productive infection of SARS-CoV-2 in ACE2+ mature enterocytes in human small intestinal enteroids. In addition to TMPRSS2, another mucosa-specific serine protease, TMPRSS4, also enhanced SARS-CoV-2 spike fusogenic activity and mediated viral entry into host cells. However, newly synthesized viruses released into the intestinal lumen were rapidly inactivated by human colonic fluids and no infectious virus was recovered from the stool specimens of COVID-19 patients. Our results highlight the intestine as a potential site of SARS-CoV-2 replication, which may contribute to local and systemic illness and overall disease progression.

## Introduction

Coronavirus disease 2019 (COVID-19) has emerged as a new world pandemic, claiming the deaths of over 160,000 people. This outbreak is caused by a novel severe acute respiratory syndrome coronavirus, SARS-CoV-2 (*1, 2*), which belongs to the family of *Coronaviridae*, a group of enveloped, non-segmented, positive-sense RNA viruses. Currently, there are no clinically approved countermeasures available for COVID-19. The SARS-CoV-2 associated disease seems to take an extra toll on the elderly, although the host immunological responses to the pathogen remain largely unknown. SARS-CoV-2 also has a more pronounced disease time course compared to some other recent respiratory viruses, raising questions about other potential modes of transmission.

The attachment of SARS-CoV-2 to the target cell is initiated by interactions between the spike glycoprotein (S) and its cognate receptor, angiotensin I converting enzyme 2 (ACE2) (*2–5*). Following receptor engagement, SARS-CoV-2 S is processed by a type II transmembrane serine protease TMPRSS2 to gain access to the host cell cytosol (*3, 6*). Interestingly, both ACE2 and TMPRSS2 are highly expressed in the gastrointestinal (GI) tract, in particular by intestinal epithelial cells (IECs), the predominant target cells for many human enteric viruses. In fact, multiple animal CoVs are natural enteric pathogens, causing GI diseases and spreading by the fecal-oral route (*7*). Notably, GI symptoms including abdominal pain and diarrhea were observed in 20-50% of COVID-19 patients and sometimes preceded the development of respiratory diseases (*8–10*). Substantial amounts of SARS-CoV-2 RNA have been consistently detected in stool specimens from COVID-19 patients (*11–15*). In a recent study, no infectious virus was successfully isolated from the feces of COVID-19 patients (*16*). However, the fecal shedding data raises the possibility that SARS-CoV-2 may act like an enteric virus and could potentially be transmitted via the fecal-oral route.

In the present study, we aimed to address: 1) do SARS-CoV-2 actively infect human IECs? 2) if so, what host factors mediate efficient replication? 3) are there replication competent viruses shed in the fecal samples of COVID-19 patients? Here, we report, for the first time, that SARS-CoV-2 apically infects human mature enterocytes and triggers epithelial cell fusion. TMPRSS2 and TMPRSS4 serine proteases mediate this process by inducing S cleavage and enhancing fusogenic activity of the virus. We also found that human colonic fluids rapidly inactivate SARS-CoV-*2 in vitro* and likely in the intestinal lumen, rendering viral RNA non-infectious in the stool specimens. Our findings have both mechanistic and translational relevance and add to our understanding of COVID-19 pathogenesis.

## Results

### SARS-CoV-2 infects human intestinal enteroids

In the intestine, ACE2 functions as a chaperone for the neutral amino acid transporter B^0^AT1 (encoded by *SLC6A19*) on IECs and regulates microbial homeostasis (*17, 18*). In fact, ACE2 expression is substantially higher in the small intestine than all the other organs including the lung in both humans and mice (Fig. S1A-B). Therefore, we sought to identify whether SARS-CoV-2 is capable of infecting IECs and understand the implications of IEC replication for fecal-oral transmission. We performed single-cell RNA-sequencing (RNA-seq) to capture the global transcriptomics in all IEC subsets in the mouse small intestinal epithelia (Fig. 1A, left panel). We found that ACE2 was predominantly expressed in Cd26^+^Epcam^+^Cd44^−^Cd45^−^ mature enterocytes (*19, 20*) (Fig. 1A, right panel). In addition, bulk RNA-seq results revealed that primary human ileum enteroids had significantly higher mRNA levels of all known CoV receptors including ACE2 than the colonic epithelial cell line HT-29 and other non-IEC human cell lines (Fig. S1C). Quantitative PCR confirmed abundant ACE2 transcript levels in both human duodenum and ileum derived enteroids (Fig. S1D). ACE2 protein was specifically co-localized with actin at the apical plasma membrane of human enteroid monolayers (Fig. 1B). Higher ACE levels correlated with more mature enterocytes present in differentiated enteroids (Fig. 1B), consistent with our scRNA-seq findings (Fig. 1A). We further confirmed in the 3D Matrigel embedded enteroids that ACE2 had an almost perfect co-localization with Villin (Fig. S1E), an IEC marker localized most strongly to the brush border of the intestinal epithelium.

**Fig. 1.**
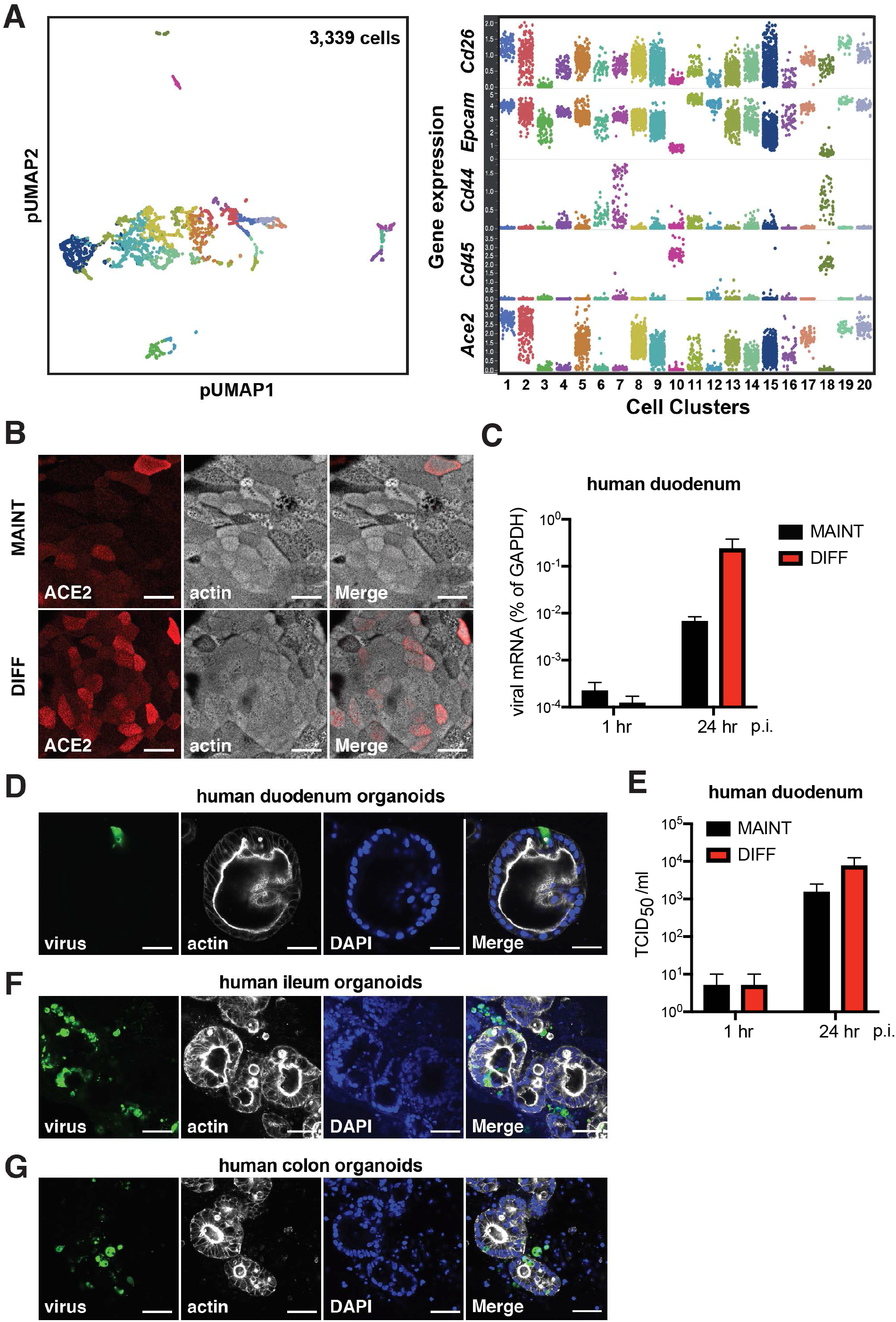
SARS-CoV-2 S chimera virus infects human small intestinal enteroids. (A) Mouse small intestinal cells were analyzed by single-cell RNA-sequencing and resolved into 20 clusters based on gene expression profiles (left panel). Transcript levels of Cd26, Epcam, Cd44, Cd45, and Ace2 were indicated for different intestinal cell subsets. Each dot represents a single cell. Note that Ace2^high^ cells are also positive for Cd26 and Epcam but negative for Cd44 and Cd45. (B) Human duodenum enteroids were cultured in the transwell monolayer system using maintenance (MAINT) or differentiation (DIFF) conditions for 3 days. Monolayers were stained for ACE2 (red) and actin (phalloidin, white). Scale bar: 32 μm. (C) Human duodenum enteroids in monolayer, cultured in either maintenance (MAINT) or differentiation (DIFF) conditions, were apically infected with 1.5×10^5^ plaque forming units (PFUs) of SARS-CoV-2 chimera virus (MOI=0.3) for 24 hours. The expression of VSV-N was measured by RT-qPCR and normalized to that of GAPDH. (D) Human duodenum enteroids in 3D Matrigel were cultured in maintenance (MAINT) media or differentiation (DIFF) media for 3 days and infected with 2.2×10^5^ PFUs of SARS-CoV-2 chimera virus for 18 hours. Enteroids were stained for virus (green), actin (phalloidin, white), and nucleus (DAPI, blue). Scale bar: 50 μm. (E) Same as (C) except that virus titers were measured using an TCID50 assay instead of viral RNA levels by QPCR. (F) Same as (D) except that human ileum enteroids were used instead. Scale bar: 80 μm. (G) Same as (D) except that human ileum enteroids were used instead. Scale bar: 80 μm.

We next took advantage of the recently established vesicular stomatitis virus (VSV)-chimera GFP reporter virus with its glycoprotein (G) genetically replaced with SARS-CoV-2 S protein (In Press). This VSV-SARS-CoV-2-S-GFP virus serves as a powerful tool to study viral entry and cell tropism. The levels of viral RNA increased by ~10,000 fold in human duodenum enteroids over 24 hours post infection (Fig. 1C), indicative of active replication. Differentiated enteroids with higher ACE2 levels (Fig. 1B) supported 39-fold higher SARS-CoV-2 chimera virus replication than stem cell based cultures (Fig. 1C). We could visualize GFP positive infected cells in duodenum enteroids in both traditional 3D (Fig. 1D) and in flipped “inside-out” models where the apical side of the IECs was on the outside of the spherical organoid (*21*) (Fig. S1F). In addition to viral RNA and protein, the amount of infectious viruses increased >1,000 fold within the intestinal epithelium at 24 hours post infection (Fig. 1E). Robust viral replication was confirmed in human ileum and colon-derived enteroids (Fig. 1F-G).

### SARS-CoV-2 actively replicates in ACE2+ human mature enterocytes

To further characterize the potential SARS-CoV-2 route of infection, we employed the 2D monolayer system (*22*), with a clear separation of apical and basal compartments. We noted that the human IECs were preferentially (> 1,000 fold) infected by the virus from the apical surface compared to the basolateral side (Fig. 2A). This data is in line with a strong apical ACE2 expression (Fig. 1B and S1E). Importantly, we found that the newly produced virus progenies were also predominantly released from the apical side into the lumen (Fig. 2B), suggesting a possibility of fecal viral shedding in COVID-19 patients.

**Fig. 2.**
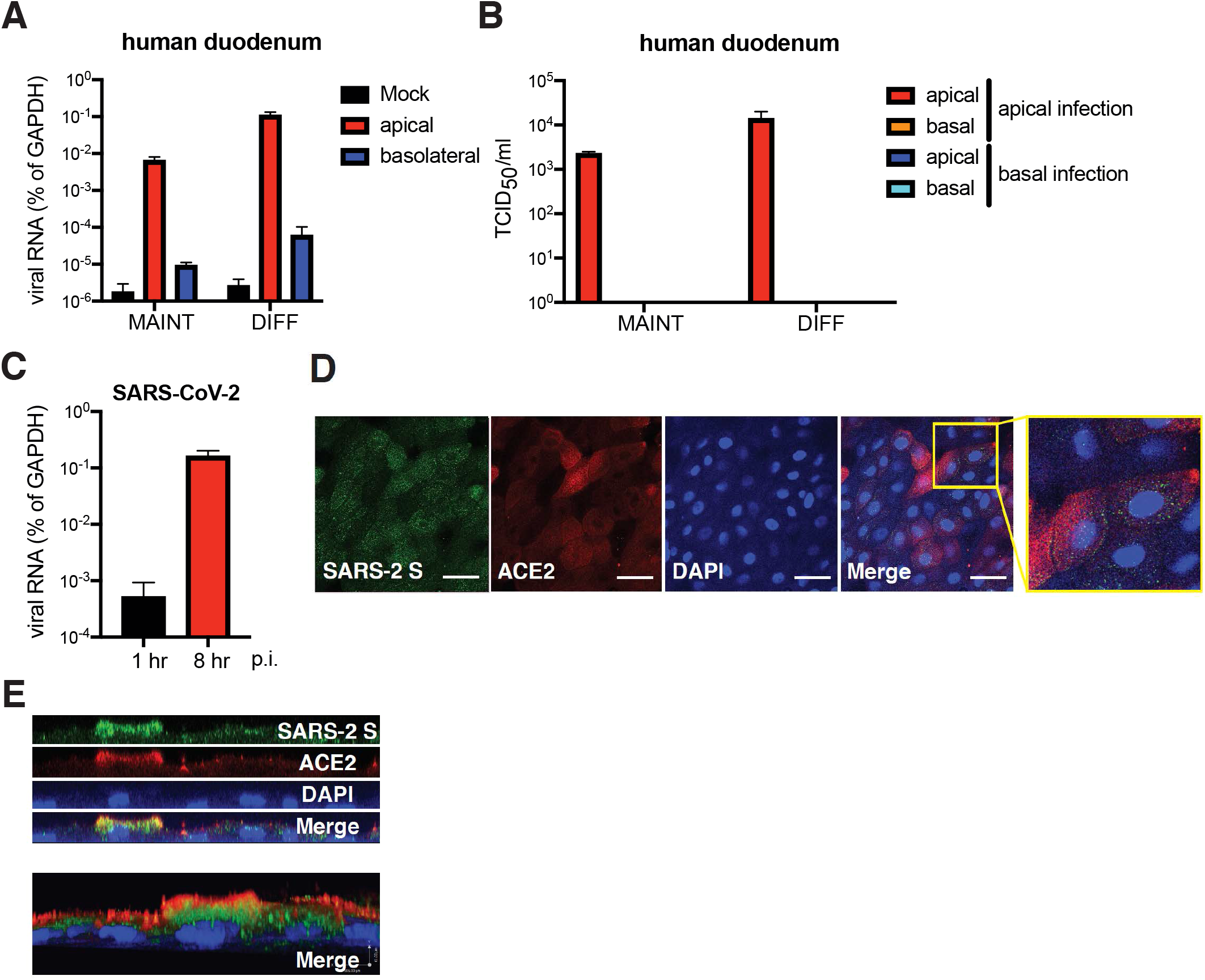
SARS-CoV-2 actively replicates in ACE2+ human mature enterocytes. (A) Human duodenum enteroids in monolayer, cultured in either maintenance (MAINT) or differentiation (DIFF) conditions, were apically or basolaterally infected with 1.5×10^5^ PFUs of SARS-CoV-2 chimera virus for 24 hours. The expression of VSV-N was measured by RT-qPCR and normalized to that of GAPDH. (B) Supernatants in both apical and basal chambers were collected from (A) and were subjected to an TCID50 assay to measure the amount of infectious viruses. (C) Differentiated duodenum enteroids in monolayer were apically infected with 2.5×10^5^ PFUs of infectious SARS-CoV-2 virus (MOI=0.5) for 8 hours. The expression of SARS-CoV-2 N was measured by RT-qPCR using a Taqman assay and normalized to that of GAPDH. (D) Same as (C) except that enteroids were fixed and stained for SARS-CoV-2 S (green), ACE2 (red), and nucleus (DAPI, blue). Scale bar: 32 μm. SARS-CoV-2 infected ACE2 positive cells are enlarged in the inset (yellow box). (E) SARS-CoV-2 infected duodenum monolayers were imaged along the z-stacks and sectioned for YZ planes (top panel) and reconstructed for 3D images (bottom panel).

To definitive pinpoint the IEC subset(s) targeted by SARS-CoV-2, we infected duodenum enteroids with the SARS-CoV-2 chimera virus and co-stained with different IEC markers. GFP signals were exclusively found in CD26 positive mature villous absorptive enterocytes and not in goblet (MUC2), enteroendocrine (CHGA), or Paneth (LYZ) cells (data not shown). Using the live infectious SARS-CoV-2 virus, we validated that viral RNA increased by 320-fold within the first 8 hours of infection (Fig. 2C). In addition, SARS-CoV-2 infection of enteroid monolayers suggested that ACE2 positive mature enterocytes were indeed the predominant target cells in the gut epithelium (Fig. 2D). Interestingly, a 3D reconstruction of confocal images showed that the S protein was highly concentrated towards the apical surface (Fig. 2E), suggesting a potential mechanism of polarized viral assembly and subsequent apical release.

While SARS-CoV induced minimal syncytia formation in the absence of exogenous proteases such as trypsin (*23, 24*), SARS-CoV-2 was able to fuse cells during natural infection of cultured cells (Fig. S2A). We also observed multiple events of syncytia formation between IECs in both 2D monolayer (Fig. S2B) and in 3D Matrigel (Fig. S2C and Video S1). The cell fusion and subsequent cytopathic effect may have important implications regarding the common GI symptoms seen in COVID-19 patients (*25–29*).

### TMPRSS2, TMPRSS4 but not ST14 mediate SARS-CoV-2 entry

Several pieces of cell line-based evidence demonstrated that the membrane-bound TMPRSS2 plays a critical role in processing S cleavage and mediating SARS-CoV-2 entry (*3, 6, 30*). Interestingly, only two other serine proteases in the same family, TMPRSS4 and matriptase (encoded by *ST14*) shared a highly specific expression pattern in human IECs (Fig. 3A). Previous studies indicated that both TMPRSS4 and ST14 facilitate influenza A virus but neither play a role in SARS-CoV infection (*31–34*). Notably, in our mouse scRNA-seq dataset, both TMPRSS4 and ST14 were found to be present in mature enterocytes and had an increased co-expression pattern with ACE2 than TMPRSS2 (Fig. S3A).

**Fig. 3.**
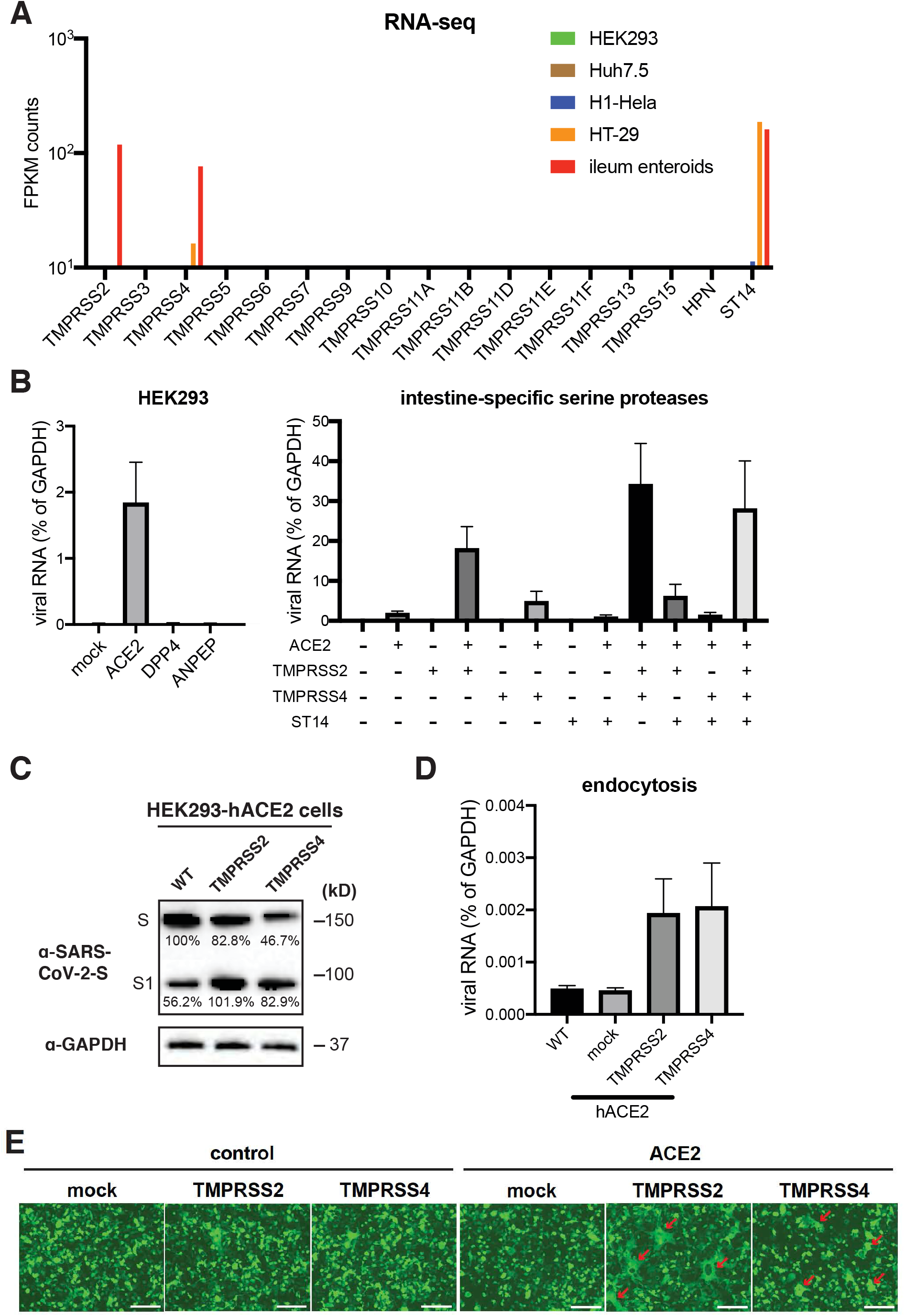
TMPRSS2, TMPRSS4 but not ST14 mediate SARS-CoV-2 entry. (A) Bulk RNA-sequencing results of intestine-specific serine protease expression in HEK293, Huh7.5, H1-Hela, HT-29 cells and human ileum enteroids. (B) HEK293 cells were transfected with pcDNA3.1-V5-ACE2, DDP4, or ANPEP for 24 hours (left panel), or transfected with indicated plasmid combination for 24 hours (right panel), and infected with 1.5×10^5^ PFUs of SARS-CoV-2 chimera virus for 24 hours. The expression of VSV-N was measured by RT-qPCR and normalized to that of GAPDH. (C) HEK293 cells stably expressing human ACE2 were transfected with C-terminally tagged SARS-CoV-2 S and TMPRSS2 or TMPRSS4 or 48 hours. The levels of S and GAPDH were measured by western blot. The intensity of bands were quantified by ImageJ and shown as percentage of the top band in lane 1. (D) HEK293 cells stably expressing human ACE2 were transfected with TMPRSS2 or TMPRSS4 for 24 hours, incubated with 5.8×10^5^ PFUs of SARS-CoV-2 chimera virus on ice for 1 hour, washed with cold PBS for 3 times, and shifted to 37 °C for another hour. The expression of VSV-N was measured by RT-qPCR and normalized to that of GAPDH. (E) Wild-type or human ACE2 expressing HEK293 cells were transfected with SARS-CoV-2 S and GFP, with or without TMPRSS2 or TMPRSS4 or 24 hours. The red arrows highlight the formation of large syncytia.

To mechanistically dissect the entry pathway of SARS-CoV-2 in IECs, we set up an HEK293 ectopic expression system to evaluate the SARS-CoV-2 chimera virus infectivity. As reported (*2, 5*), ACE2 conferred permissiveness to SARS-CoV-2 infection (Fig. 3B and S3B). While TMPRSS2 alone did not mediate viral infection, co-expression of TMPRSS2 significantly enhanced ACE2-mediated infectivity (Fig. 3B). Importantly, we found that expression of TMPRSS4 but not ST14 also resulted in a significant increase in the levels of viral RNA and infectious virus titers in the presence of ACE2 (Fig. 3B and S3C). TMPRSS4 and TMPRSS2 had an additive effect and mediated the maximal infectivity in cell culture (Fig. 3B and S3C-E).

We reasoned that TMPRSS4 may function as a cell surface serine protease that enhances S cleavage to promote viral entry. To test this hypothesis, we co-expressed a C-terminally Strep-tagged full-length SARS-CoV-2 S protein in an HEK293 cell line that stably expresses ACE2 with or without additional introduction of TMPRSS2 or TMPRSS4. In mock cells, we readily observed the full-length S and a cleaved product that corresponded to the size of S1 fragment, presumably cleaved by furin protease that is ubiquitously expressed (*35*). Importantly, the expression of TMPRSS2 or TMPRSS4 enhanced S cleavage, as evidenced by the reduction of full-length S and increase of S2 levels (Fig. 3C).

Based on these results, we hypothesize that TMPRSS serine proteases assist virus infection by inducing S cleavage and exposing the fusion peptide for efficient viral entry. Indeed, TMPRSS4 expression did not affect virus binding (data not shown) whereas the endocytosis was significantly enhanced (Fig. 3D). To examine S protein’s fusogenic activity, we examined SARS-CoV-2 S mediated cell-cell fusion in the presence of absence of TMPRSS2 or TMPRSS4. Previous work with SARS-CoV and MERS-CoV suggested S mediated membrane fusion takes place in a cell-type dependent manner (*36*). We found that the ectopic expression of S alone was sufficient to induce syncytia formation, independent of the virus infection (Fig. 3E). This process was dependent on a concerted effort from ACE2 and TMPRSS serine proteases. TMPRSS4 expression triggered S-mediated cell-cell fusion, although to a lesser extent than TMPRSS2 (Fig. 3E). Collectively, we have shown that TMPRSS2 and TMPRSS4 activate SARS-CoV-2 S and enhance membrane fusion and viral endocytosis into host cells.

### TMPRSS2 and TMPRSS4 promote SARS-CoV-2 infection in enteroids

To further probe the TMPRSS mechanism of action and physiological relevance and to recapitulate the IEC gene expression pattern *in vivo* (Fig. S3A), we designed an *in vitro* co-culture system where ACE2 and S were expressed on target and donor cells respectively. Based on the scRNA-seq data (Fig. S3A), we constructed an HEK293 stable cell line that co-expresses ACE2 and TMPRSS4 to mimic mature enterocytes and another HEK293 cell line that stably expresses TMPRSS2 to mimic goblet or other secretory IEC types (Fig. 4A, left panel). Target cells were transfected with GFP and mixed at 1:1 ratio with SARS-CoV-2 S containing donor cells that were transfected with TdTomato. We found that TMPRSS4 was able to function *in cis*, i.e. on the same cells as ACE2, and induced fusion of GFP positive cells (Fig. 4A, right panel), as was shown in Fig. 3E. Importantly, we discovered that TMPRSS2 could act *in trans* and its expression on adjacent cells markedly promoted the formation of stronger and larger cell-cell fusion (Fig. 4A, right panel).

**Fig. 4.**
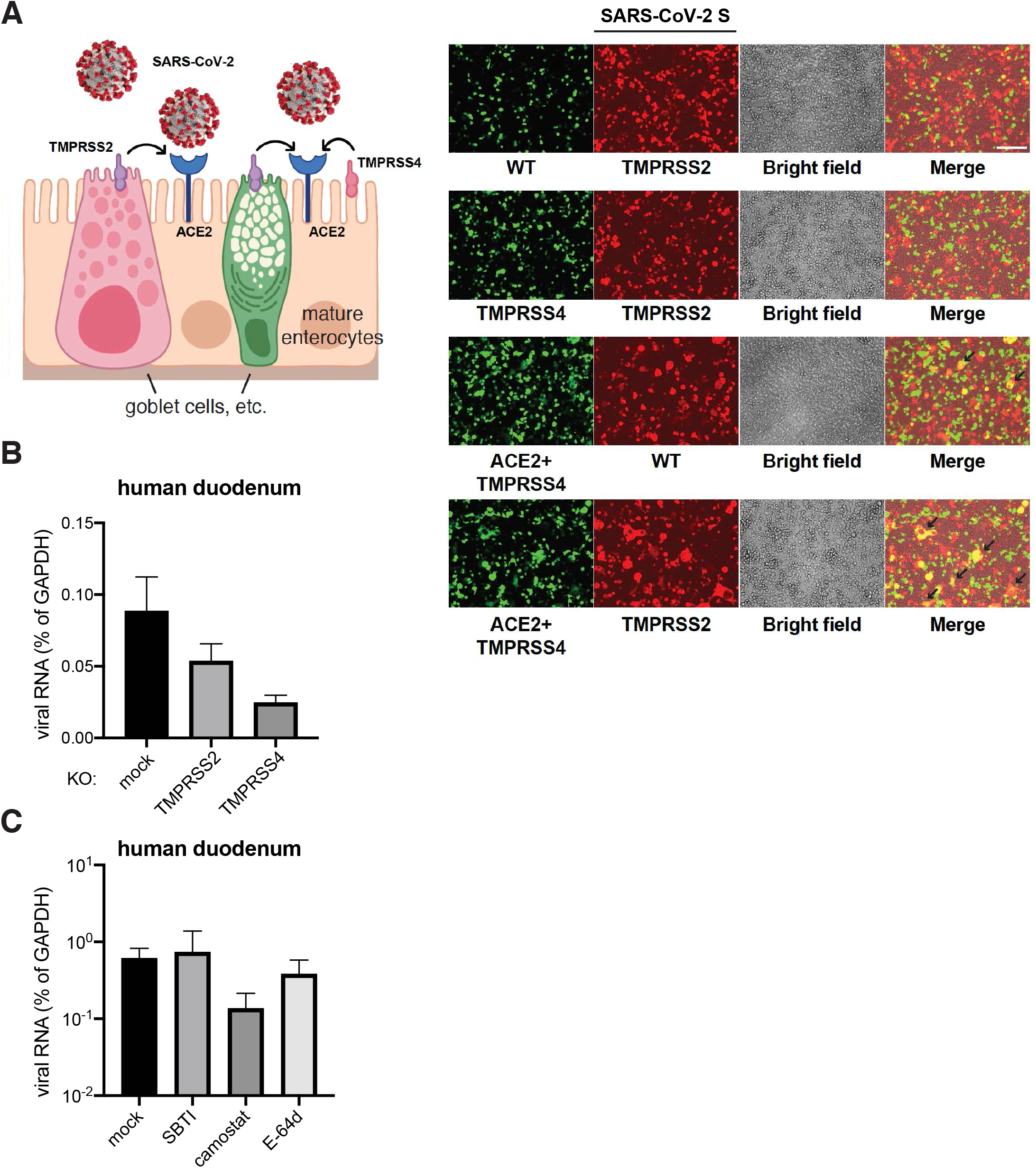
TMPRSS2 and TMPRSS4 promote SARS-CoV-2 infection in enteroids. (A) Schematic diagram of SARS-CoV-2 infection of human mature enterocytes (left panel). An HEK293 stable cell line expressing ACE2 and TMPRSS4 were transfected with GFP and an HEK293 stable cell line expressing TMPRSS2 were transfected with SARS-CoV-2 S and TdTomato for 24 hours. These two cell lines were then mixed at 1:1 ratio and cultured for another 24 hours. Note the formation of cell-cell fusion (yellow), highlighted by black arrows. (B) Human duodenum enteroids in 3D Matrigel were transduced with lentiviruses encoding Cas9 and sgRNA against TMPRSS2 or TMPRSS4 (oligonucleotide information in Table S1). Gene knockout enteroids were seeded into monolayers and infected with 1.5×10^5^ PFUs of SARS-CoV-2 chimera virus for 24 hours. The expression of VSV-N was measured by RT-qPCR and normalized to that of GAPDH. (C) Human duodenum enteroids seeded into collagen-coated 96-well plates were differentiated for 3 days, pre-treated with 50 μg/ml of soybean trypsin inhibitor (SBTI), 10 μM of camostat mesylate, or 10 μM of E-64d for 30 minutes, and infected with 1.5×10^5^ PFUs of SARS-CoV-2 chimera virus for 24 hours. The expression of VSV-N was measured by RT-qPCR and normalized to that of GAPDH.

Next, to determine the function of TMPRSS serine proteases in facilitating SARS-CoV-2 infection of primary human IECs, we used a CRISPR/Cas9-based method to genetically delete TMPRSS2 or TMPRSS4 in human duodenum enteroids. Efficient knockout was confirmed by western blot (Fig. S4A). Importantly, abrogating TMPRSS4 expression led to a 4-fold reduction in SARS-CoV-2 chimera virus replication in human enteroid, even more significant than TMPRSS2 knockout (Fig. 4B), highlighting its importance in mediating virus replication in primary cells.

In parallel to genetic depletion, we also tested the effect of pharmacological inhibition of TMPRSS serine proteases on virus replication. We pre-treated enteroids with camostat mesylate, a selective inhibitor of TMPRSS over other serine proteases including trypsin, prostasin and matriptase (*37*). While camostat treatment significantly inhibited SARS-CoV-2 chimera virus infection, soybean trypsin inhibitor (SBTI) and E-64d, a cysteine protease inhibitor that blocks cathepsin activity and has excellent inhibitory activity against SARS-CoV-2 *in vitro (3)*, did not had a major impact on virus replication in enteroids (Fig. 4C).

Taken together, our data support a model where SARS-CoV-2, at least in human IECs, seems to be more reliant on the membrane-bound TMPRSS serine proteases (Fig. 4A, left panel), as compared to SARS-CoV and other CoVs that can efficiently utilize both TMPRSS2 and endosome-localized cathepsins as alternative routes of entry (*38, 39*).

### SARS-CoV-2 is rapidly inactivated in the human GI tract

Our current results strongly suggest that SARS-CoV-2 viruses are able to infect human IECs apically and are released via the apical route into the lumen (Fig. 2). Human enteric viruses that spread via the fecal-oral route typically withstand the harsh environment in the GI tract, including the low pH of gastric fluids, bile and digestive enzymes in the small intestine, dehydration and exposure to multiple bacterial by-products in the colon. We sought to investigate the stability of SARS-CoV and SARS-CoV-2 chimera viruses in different human gastric and intestinal fluids. Compared to rotavirus, a prototypic human enteric virus known to successfully transmit fecal-orally (*40*), both CoV chimera viruses quickly lost infectivity in a low pH condition (Fig. S5A). However, SARS-CoV-2-S exhibited greater stability than SARS-CoV-S in human small intestinal fluids that contain biological surfactants including taurocholic acid sodium salt and lecithin (Fig. 5A). Interestingly, SARS-CoV-2-S chimera virus was not resistant to certain components in the human colonic fluids (Fig. 5A). The virus titers decreased by 100-fold within 1 hour and no infectious virus was detectable at the 24 hour time point (Fig. 5A). In contrast, rotavirus remained stable in all gastric and enteric fluids tested (Fig. S5A-B).

**Fig. 5.**
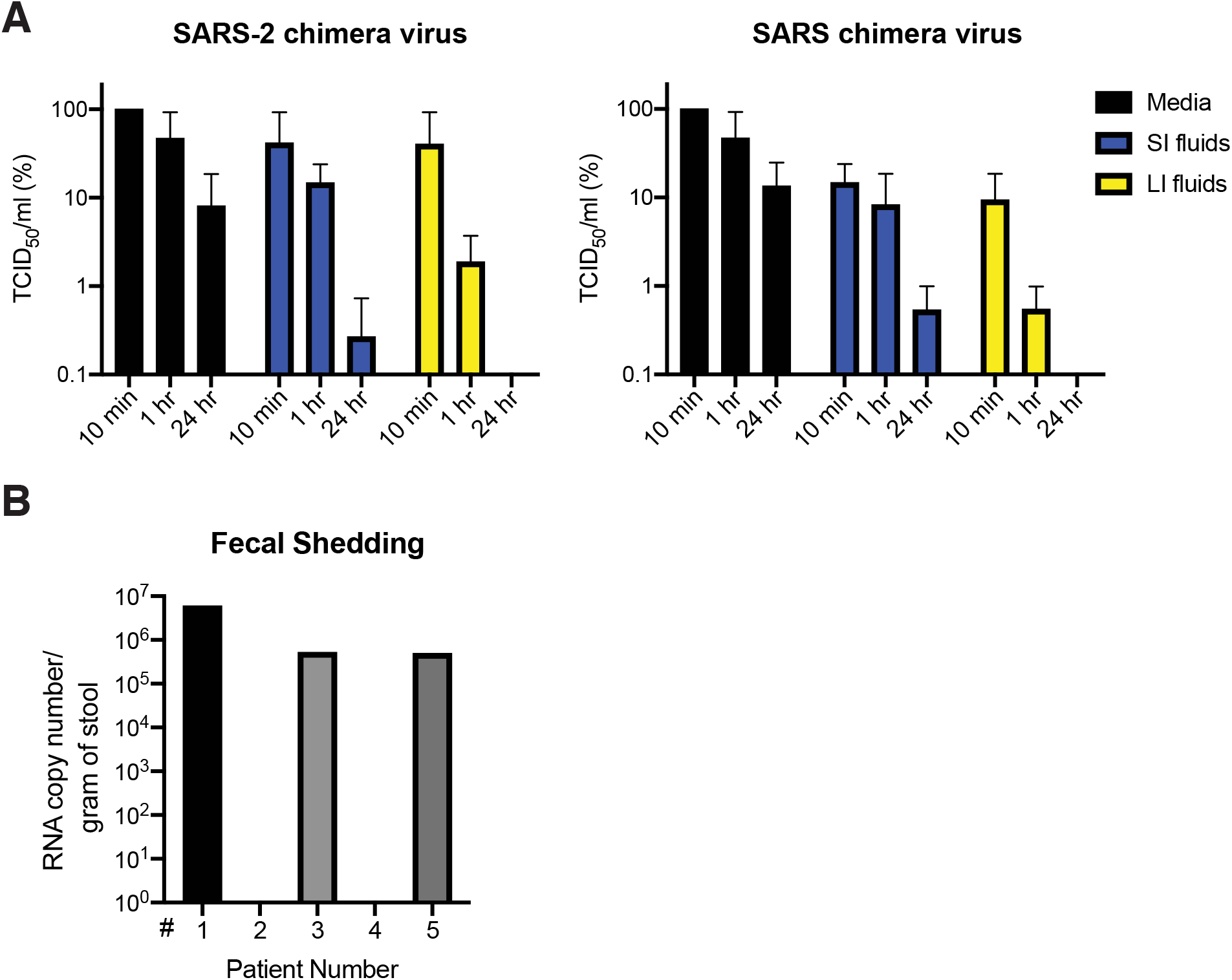
SARS-CoV-2 rapidly lose infectivity in the human GI tract. (A) 2.9×10^5^ PFUs of SARS-CoV-2 and SARS-CoV chimera viruses were incubated with M199 media, human small intestinal (SI) fluids, or human large intestinal (LI) fluids for indicated time points at 37 °C. The infectivity of viruses was subsequently determined by a standard TCID50 cell culture based assay. (B) Stool specimens from five COVID-19 patients were collected and subjected to QPCR experiments to quantify the absolute levels of SARS-CoV-2 N gene.

Combined with the human enteroid results (Fig. 1-2), we hypothesize that the SARS-CoV-2 has the potential to and likely does replicate in human IECs but then get quickly inactivated the GI tract. We collected stool specimens from a small group of COVID-19 patients. From 3 out of 5 fecal samples, we detected high RNA copy numbers of SARS-CoV-2 viral genome (Fig. 5B). However, we were unable to recover any infectious virus using a highly sensitive cell-based assay (*6*).

## Discussion

In this study we set out to address an important basic and clinically relevant question: Is the persistent viral RNA seen in COVID-19 patients’ stool infectious and transmissible? We showed that despite SARS-CoV-2’s ability to establish robust infection and replication in human IECs (Fig. 1-2), the virus is rapidly inactivated by colonic fluids and we were unable to detect infectious virus in fecal samples (Fig. 5). Thus, the large quantities of viral RNA that transit through the GI tract and shed into the feces are likely not infectious in nature. Due to the limitation of sample size, we cannot definitely conclude that fecal-oral transmission of COVID-19 does not exist. However, our data is consistent with the previous reports from SARS-CoV and MERS-CoV publications which concluded that despite long duration of viral RNA shedding, no infectious virus could be recovered from the patients’ feces (*41, 42*).

Intriguingly, the small intestinal fluids, which contain taurocholic acid sodium salt and lecithin, important surfactant components of the bile, did not inactivate virus as expected (Fig. 5A). In addition, we also observed that the virus titers did not drop in the presence of 0.03% bovine bile (data not shown), the highest concentration that did not induce cytotoxicity, contradicting the known fragility of lipid bilayers and sensitivity to bile salts which serve as detergents in gut. Recent studies showed a heavy glycosylation of SARS-CoV-2 S protein (*43*) and it is possible the sugar coating conferred some stability against the enzymatic digestion and bile salt solubilization. Two recent publications did demonstrate that SARS-CoV-2 has unique stability in the environment, at least in a laboratory setting (*44, 45*). Further work is needed to determine which bioactive(s) in the colonic fluids that are absent from small intestinal fluids function to potentially disrupt viral lipid membranes *in vivo*.

Mechanistically, recent work from multiple groups showed a key role of TMPRSS2 in cleaving SARS-CoV-2 S protein. Here, we provide data that besides TMPRSS2, TMPRSS4 also increases SARS-CoV-2 infectivity, at least in gut epithelial cells (Fig. 3-4). We observed an intriguing additive effect of the two enzymes, as both target single arginine or lysine residues (R/K↓). One may speculate that this synergy stems from a distinct subcellular localization in neighboring cells (Fig. 4A). Future work with recombinant proteins and enzymatic digestion may help address whether S cleavage is sequential and whether the two enzymes act on different sites on the S protein.

Finally, we observed strong syncytia formation in human enteroids (Fig. S2B-C). These results may have great implications for virus cell-cell spread and evasion from antibody neutralization (*46*). It is unclear at this point whether the fused IECs will still undergo anoikis or an alternative accelerated cell death, which may cause a breach in barrier integrity and give rise to GI pathology seen in COVID-19 patients. It is possible that in the small intestine, whereas SARS-CoV-2 is relatively stable, additional proteases such as trypsin likely enhance viral pathogenesis by triggering more robust IEC fusion (Fig. S2A). In this sense, although viruses are not released into the basolateral compartment (Fig. 2B), one can imagine that a leaky gut will allow the virus to disseminate to other systemic organs including the lung and liver. This hypothesis is consistent with the clinical observation that in some COVID-19 patients, the GI symptoms preceded the respiratory illness (*8, 9*). Whether the gut serves as a primary site of infection and whether it contributes to systemic diseases in individual patients require further studies using appropriate animal models.

## Materials and Methods

### Plasmids, Cells, Reagents, and Viruses

#### Plasmids

Human ACE2, DPP4, ANPEP, TMPRSS2, TMPRSS4, and ST14 were cloned into a pcDNA3.1/nV5-DEST vector with an N-terminal V5 tag and a neomycin selection marker. Human TMPRSS2 and TMPRSS4 were also cloned into a pLX304 lentiviral vector with a C-terminal V5 tag and a blasticidin selection marker. pEGFP-N1 and pCMV-TdTomato were commercially purchased from Clontech. Codon-optimized SARS-CoV-2 S was a kind gift from Dr. Nevan J. Krogan at UCSF (*47*).

#### Cells

African Green Monkey kidney epithelial cell lines MA104 (CRL-2378.1) and human embryonic kidney cell line HEK293 (CRL-1573) were originally obtained from American Type Culture Collection (ATCC) and cultured in complete M199 medium and complete DMEM medium, respectively. HEK293 cells stably expressing human ACE2 or TMPRSS2 were selected under 500 μg/ml G418. HEK293 cells stably expressing human ACE2 and TMPRSS2 were selected under 500 μg/ml G418 and 5 μg/ml blasticidin.

#### Reagents

Human gastric fluids (Fasted State Simulated Gastric Fluid/FaSSGF, pH 1.6), small intestinal fluids (Fasted State Simulated Intestinal Fluid/FaSSIF-V2, pH 6.5), colonic fluids (Fasted State Simulated Colonic Fluid/FaSSCoF, pH 7.8) were purchased from BioRelevant, UK and reconstituted based on the manufacturer’s instructions. Soybean trypsin inhibitor (SBTI), camostat mesylate, and E-64d were purchased from Selleckchem. Primary antibodies used in this study included: GAPDH (631402, Biolegend); GFP (2555S, Cell Signaling); SARS-CoV-2-S (S1) (PA5-81795, Thermo Fisher); TMPRSS2 (sc-515727, Santa Cruz); TMPRSS4 (sc-376415, Santa Cruz); and V5 (13202S, Cell Signaling).

#### Viruses

Human rotavirus WI61 strain was propagated as described before (*48*). VSV-SARS-CoV-2-S-GFP and VSV-SARS-CoV-S-GFP were propagated in MA104 cells in T175 flasks and virus sequences can be found in Table S1. Plaque assays were performed in MA104 cells seeded in 6-well plates using an adapted version of the rotavirus plaque assay protocol (*19*). TCID50 assays were performed in MA104 cells seeded in flat-bottom 96-well plates using serial dilutions. Virus infections were performed with the initial inoculum removed at 1 hour post adsorption.

### Human intestinal enteroids

One duodenum (#CD94), five ileum (#14-75, #211D, #262D-2, #265D, #251D-2), and one colon (#235A) enteroids were used in this study. All enteroids were derived from de-identified tissue from healthy, non-IBD subjects undergoing colonoscopy and provided informed consent at Stanford University and Washington University in St. Louis. In brief, enteroids were cultured in maintenance media (advanced DMEM/F12 media supplemented with L-WRN-conditioned media that contains Wnt-3a, R-spondin 3, and Noggin) in 3D Matrigel in 24-well plates (*49*). For passing enteroids, Y27632 and CHIR99021 were added. For differentiation, conditioned media was replaced with 50% Noggin in the form of recombinant proteins. Enteroids, when required, were digested into single cells using TrypLE and seeded into collagen-coated transwells (0.33cm^2^, 0.4μm pore size, Polycarbonate, Fisher #07-200-147) in ROCK Inhibitor Y-27632 supplemented maintenance media in 24-well plates. TEER measurement was performed using the Millicell ERS-2 Voltohmmeter (Millipore). Enteroids in monolayers were used for virus infection if the cultures had TEER greater than 1000 Ω.cm2 at 5 days post seeding. For CRISPR/Cas9 knockout, enteroids in 3D Matrigel were digested with 5 mM EDTA and transduced with lentiviral vectors encoding Cas9 and single-guide RNA against TMPRSS2 or TMPRSS4 (see Table S1) in the presence of polybrene (8 μg/ml). At 48 hours post transduction, puromycin (2 μg/ml) was added to the maintenance media. Puromycin was adjusted to 1 μg/ml upon the death of untransduced control enteroids.

### RNA extraction and quantitative PCR

Total RNA was extracted from cells using RNeasy Mini kit (Qiagen) and reverse transcription was performed with High Capacity RT kit and random hexamers as previously described (*50*). QPCR was performed using the AriaMX (Agilent) with a 25 μl reaction, composed of 50 ng of cDNA, 12.5 μl of Power SYBR Green master mix (Applied Biosystems), and 200 nM both forward and reverse primers. All SYBR Green primers and Taqman probes used in this study are listed in Table S1.

### Bright-field and immunofluorescence microscopy

For brightfield and epifluorescence, cultured cells or human enteroids were cultured in 24-well plates and images were taken by REVOLVE4 microscope (ECHO) with a 10X objective. For confocal microscopy, enteroids were seeded into 8-well Nunc chamber slides (Thermo Fisher), cultured under maintenance or differentiation conditioned, infected with SARS-CoV-2 chimera virus, and fixed with 4% paraformaldehyde as previously described (*48*). Samples were then stained with the following primary antibodies or fluorescent dyes: ACE2 (sc-390851 AF594, Santa Cruz), DAPI (P36962, Thermo Fisher), SARS-CoV-2 S (CR3022 human monoclonal antibody (*51*)), Villin (sc-58897 AF488, Santa Cruz), and phalloidin (Alexa 647-conjugated, Thermo Fisher). Stained cells were washed with PBS, whole mounted with Antifade Mountant, and imaged with Zeiss LSM880 Confocal Microscope at the Molecular Microbiology imaging core facility at Washington University in St. Louis. Z-stack was applied for imaging 3D human enteroids. Images were analyzed by Volocity v6.3 (PerkinElmer) and quantification was determined by CellProfiler (Broad Institute).

### Single-cell RNA-sequencing analysis

Small intestinal IEC isolation by EDTA-DTT extraction method and sorting were performed as previously described (*19, 52*). Cells were stained with LIVE/DEAD Aqua Dead Cell Stain Kit (Thermo Fisher) and live cells were sorted using a BD FACSAria II flow cytometer. Preparation of single cell suspensions was performed at the Stanford Genome Sequencing Service Center (GSSC). Cell suspensions were processed for single-cell RNA-sequencing using Chromium Single Cell 3’ Library and Gel Bead Kit v2 (10X Genomics, PN-120237) according to 10X Genomics guidelines. Libraries were sequenced on an Illumnia NextSeq 500 using 150 cycles high output V2 kit (Read 1-26, Read2-98 and Index 1-8 bases). The Cell Ranger package (v3.0.2) was used to align high quality reads to the mouse genome (mm10). Normalized log expression values were calculated using the scran package (*53*). Highly variable genes were identified using the FindVariableGenes function (Seurat, v2.1) as described (*54*). Batch effects from technical replicates were removed using the MNN algorithm as implemented in the batchelor package’s (v1.0.1) fastMNN function. Imputed expression values were calculated using a customized implementation (https://github.com/kbrulois/magicBatch) of the MAGIC (Markov Affinity-based Graph Imputation of Cells) algorithm (*55*) and optimized parameters (t = 2, k = 9, ka = 3). Cells were classified as dividing or resting using a pooled expression value for cell cycle genes (Satija Lab Website: regev_lab_cell_cycle_genes). For UMAP and tSpace embeddings, cell cycle effects were removed by splitting the data into dividing and resting cells and using the fastMNN function to align the dividing cells with their resting counterparts. Dimensionality reduction was performed using the UMAP algorithm and nearest neighbor alignments for trajectory inference and vascular modeling were calculated using the tSpace algorithm. Pooled expression of all viral genes was calculated using the AddModuleScore function from the Seurat package.

## Acknowledgements

We really appreciate the assistance from Matthew Williams (Molecular Microbiology Media and Glassware Facility), Wandy Betty (Molecular Microbiology Imaging Facility), Roseanne Zhao (Division of Rheumatology) and Philip Mudd (Division of Emergency Medicine) for fecal specimen collection. SARS-CoV-2 Taqman probe and viral RNA standards were prepared by Adam Bailey (Division of Infectious Diseases). We thank Krista Maria Hennig and Peter Maguire from the Stanford Genome Sequencing Service Center for their help with the single-cell RNA-seq experiments.

---

**Fig. S1.**
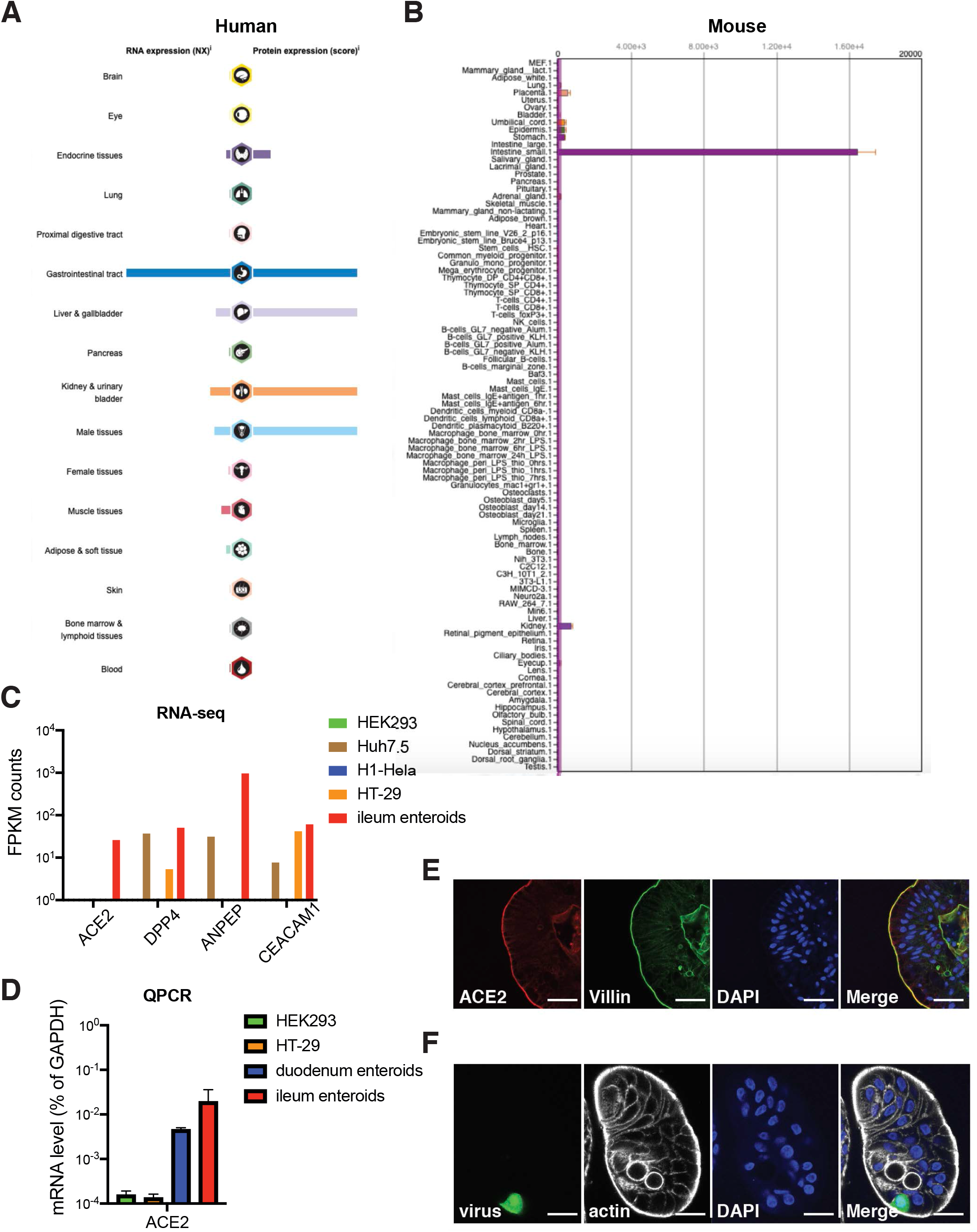
ACE2 is highly expressed in the human intestine. (A) ACE2 expression in different human tissues and organs from human protein atlas (www.proteinatlas.org/). (B) Ace2 expression in different mouse tissues and cell types from BioGPS (biogps.org). (C) Bulk RNA-sequencing results of ACE2 and other CoV receptor expression in HEK293, Huh7.5, H1-Hela, HT-29 cells and human ileum enteroids. (D) ACE2 expression level in HEK293, HT-29 cells, duodenum enteroids and ileum enteroids were measured by RT-qPCR and normalized to that of GAPDH. (E) Human duodenum enteroids in 3D Matrigel were cultured in differentiation media for 3 days and stained for ACE2 (red), villin (green), and nucleus (DAPI, blue). Scale bar: 50 μm. (F) Differentiated human duodenum enteroids were allowed to flip inside out, infected with 1.5×10^5^ PFUs of SARS-CoV-2 chimera virus for 24 hours. Enteroids were stained for virus (green), actin (phalloidin, grey), and nucleus (DAPI, blue). Scale bar: 32 μm.

**Fig. S2.**
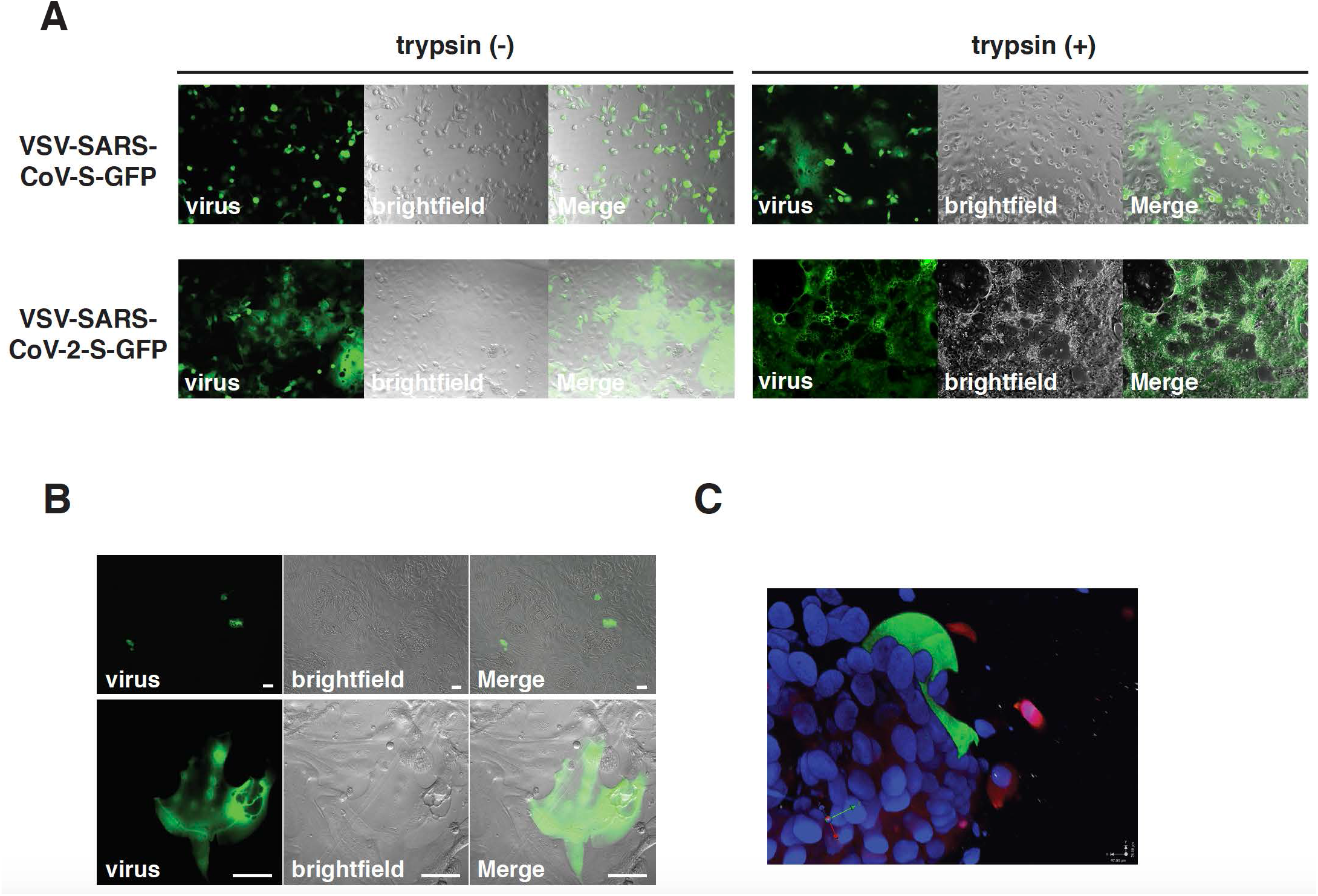
SARS-CoV-2 chimera virus induces syncytia formation in cell culture and human intestinal cells. (A) MA104 cells in serum-free media were infected with SARS-CoV and SARS-CoV-2 chimera viruses (MOI=0.1) with or without trypsin (0.5 μg/ml) for 48 hours. (B) Human ileum enteroids seeded into collagen-coated 96-well plates were apically infected with SARS-CoV-2 chimera virus (MOI=0.1) for 24 hours. (C) Human ileum enteroids in 3D Matrigel were infected with SARS-CoV-2 chimera virus (MOI=0.1) for 24 hours and stained for virus (green), ACE2 (red), and nucleus (DAPI, blue). Also see Supplementary Video 1 for 3D view.

**Fig. S3.**
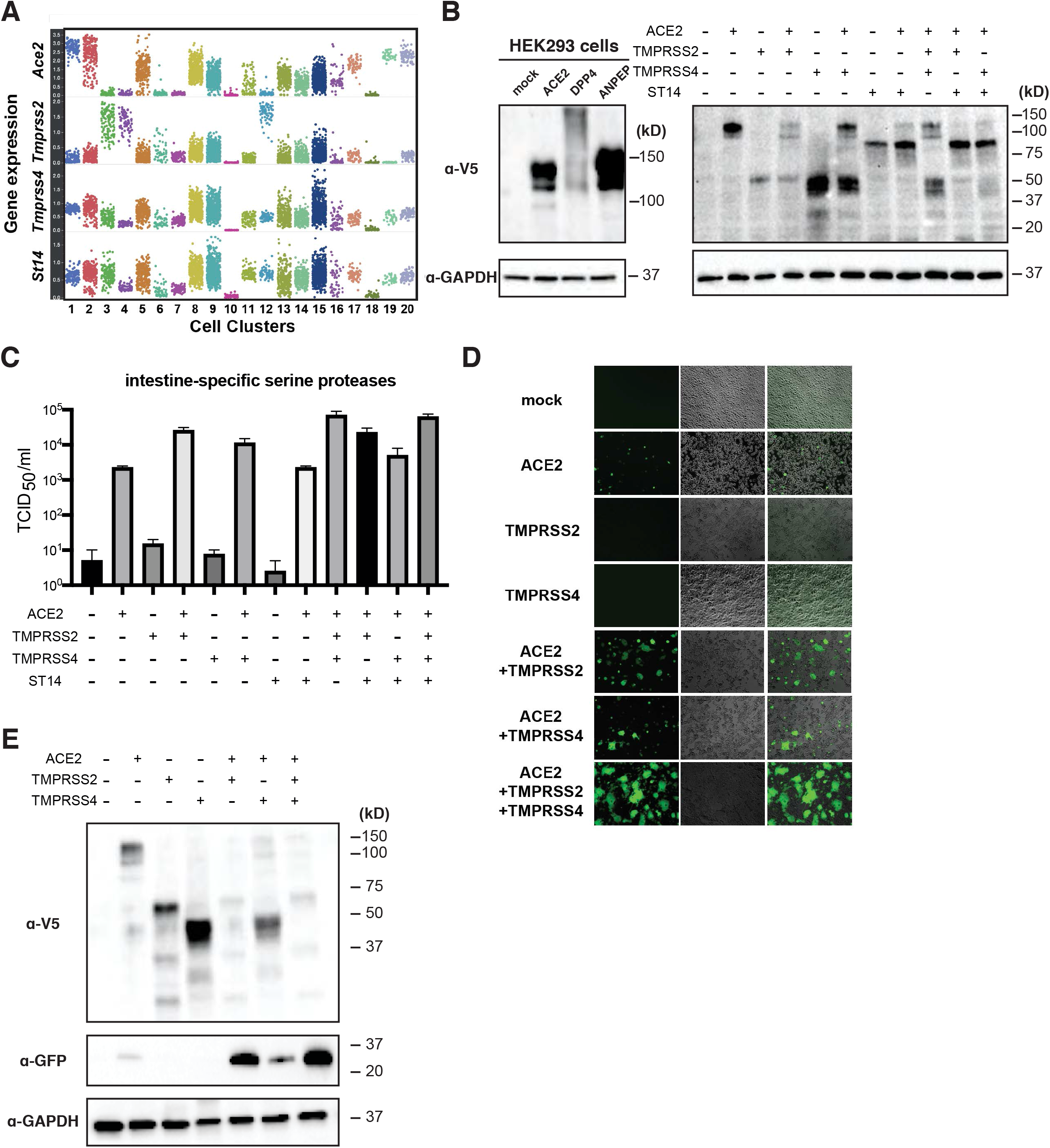
TMPRSS2 and TMPRSS4 enhance SARS-CoV-2 infectivity. (A) Transcript levels of Ace2, Tmprss2, Tmprss4, and St14 were indicated for different intestinal cell subsets. Each dot represents a single cell. (B) HEK293 cells were transfected with pcDNA3.1-V5-ACE2, DDP4, or ANPEP for 24 hours (left panel), or transfected with indicated plasmid combination for 24 hours (right panel). The levels of V5 and GAPDH were measured by western blot. (C) HEK293 cells were transfected with indicated plasmid combination for 24 hours (right panel), and infected with 1.5×10^5^ PFUs of SARS-CoV-2 chimera virus for 24 hours. The amount of infectious viruses was measured using an TCID50 assay. (D) Same as (C) except that virus-infected cells (GFP) were imaged. (E) Same as (C) except that the levels of V5, GFP, and GAPDH were measured by western blot.

**Fig. S4.**
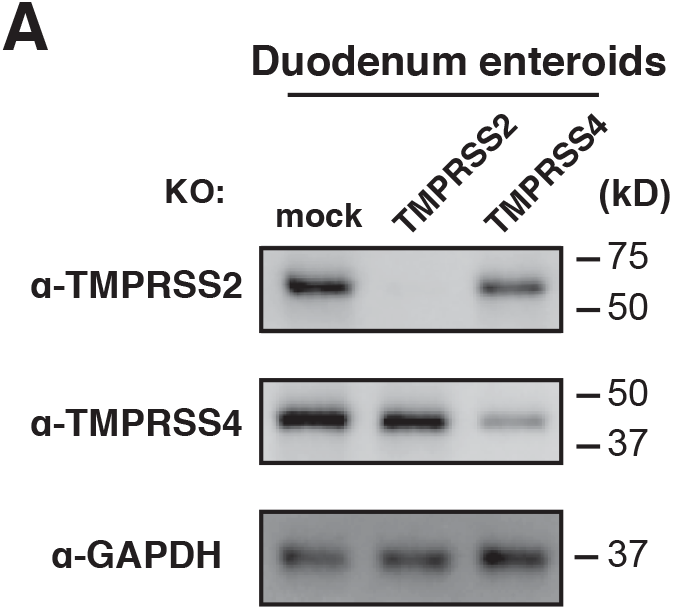
Validation of CRISPR/Cas9 mediated knockout of TMPRSS2 and TMPRSS4 in human enteroids. (A) Human duodenum enteroids were transduced with lentiviral vectors encoding Cas9 and single-guide RNA targeting TMPRSS2 or TMPRSS4, selected under puromycin (2 μg/ml) for 7 days, allowed for expansion, and harvested for western blot examining the protein levels of TMPRSS2, TMPRSS4 and GAPDH.

**Fig. S5.**
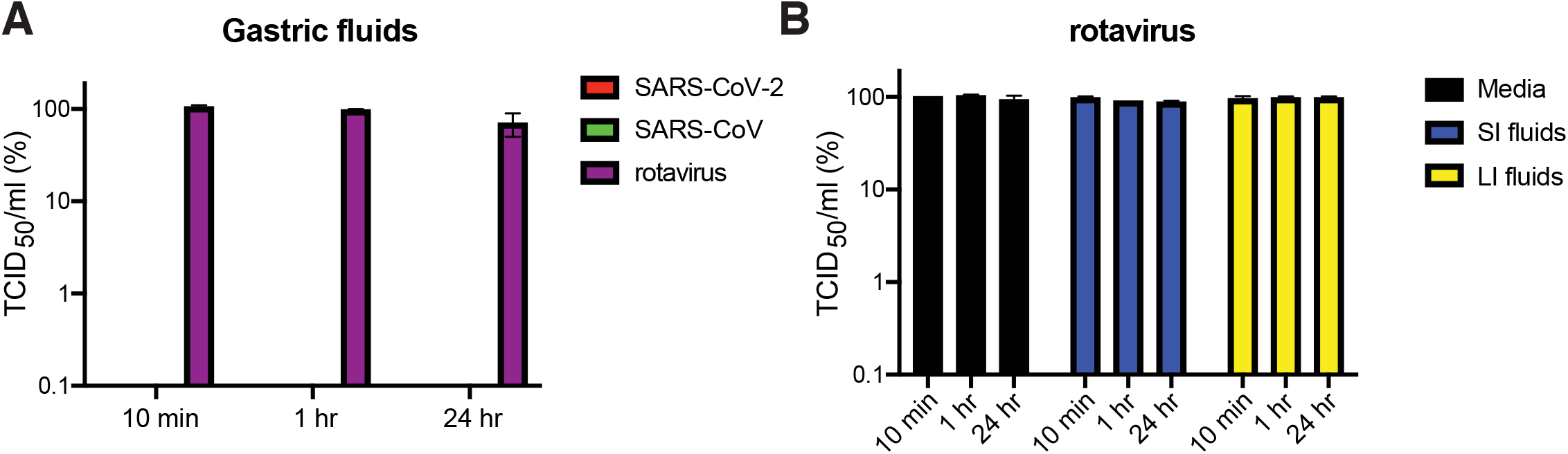
Rotavirus remains infectious in human gastric and intestinal fluids. (A) 2.9×10^5^ PFUs of SARS-CoV-2, SARS-CoV chimera viruses and rotavirus were incubated with M199 media or human gastric fluids for indicated time points. The amount of infectious viruses was determined by a TCID50 assay. (B) Same as (A) except that rotavirus and human small intestinal (SI) and large intestinal (LI) fluids were used instead.

Video S1. Formation of syncytia in 3D human duodenum enteroids infected by SARS-CoV-2 GFP chimera virus

Table S1. QPCR and CRISPR deletion oligonucleotide information

